# Differential regulation of mRNA stability modulates transcriptional memory and facilitates environmental adaptation

**DOI:** 10.1101/2022.05.05.490809

**Authors:** Bingnan Li, Patrice Zeis, Alisa Alekseenko, Gen Lin, Manu M Tekkedil, Lars M. Steinmetz, Vicent Pelechano

## Abstract

Transcriptional memory, by which cells respond faster to repeated stimuli, is key for cellular adaptation and organism survival. Factors related to chromatin organization and activation of transcription have been shown to play a role in the faster response of those cells previously exposed to a stimulus (primed). However, the contribution of post-transcriptional regulation is not yet explored. Here, combining flow cytometry and high throughput sequencing, we perform a genome-wide screen to identify novel factors modulating transcriptional memory in *S. cerevisiae* in response to galactose nutrition sources. In addition to the well-known chromatin factors modulating transcriptional memory, we find that depletion of the nuclear RNA exosome increases *GAL1* expression in primed cells. We perform a genome-wide characterisation of this process and show that changes in nuclear surveillance factor association can enhance both gene induction and repression in primed cells. Finally, we show that in addition to nuclear mRNA degradation, differences in cytoplasmic mRNA decay also modulate transcriptional memory and contribute to faster gene expression remodelling in primed cells. Our results demonstrate that mRNA post-transcriptional regulation, and not only transcription regulation, should be considered when investigating gene expression memory.

## Introduction

Transcriptional memory is the process by which previously expressed genes remain poised for faster reactivation^12^. The ability to “remember” previous stimuli and respond faster to future events is key for cellular adaptation and organism survival. For example, in *Saccharomyces cerevisiae* it is well known that previous exposure to different carbon sources^3–5^ or stress conditions^6^ facilitates future gene expression changes. A well-studied case in budding yeast is the galactose-induced transcriptional memory that leads to faster reactivation of the yeast *GAL* genes^4,7,8^. Transcriptional memory also plays a role in environmental stress adaptation in plants ^910^ and facilitates faster interferon-γ response in humans^11^. In the disease context, transcriptional memory contributes to trained immunity in macrophages, where previous exposure to LPS or ß-glucan can modulate immune tolerance via epigenetic reprogramming^12,13^, and also explains how prior inflammation can modulate tissue regeneration^14^. Thus, the ability of cells to alter future gene expression based on previous stimuli is a fundamental phenomenon in biology and key to understanding cell identity and disease progression.

Multiple mechanisms have been implicated in transcriptional memory, most of which act at the chromatin level (reviewed in ^12^). Mechanisms associated with transcriptional memory include chromatin remodelling^7^, incorporation of the histone variant H2A.Z or presence of H3K4me2^15,16,17^, association to the nuclear pore, DNA topology^4,15,18^ or even long noncoding RNA (lncRNA)-based formation of R-loops^19^. Although we know that many factors are involved in transcriptional memory, our knowledge of this process is far from complete. This is especially true regarding non-chromatin-related factors modulating transcriptional memory. As an example of a factor independent of chromatin organization, the cytoplasmic accumulation of *GAL1* (galactokinase 1) protein can facilitate faster response in galactose in primed yeast cells by acting as a positive transcriptional regulator of *GAL* genes^20,21^.

To identify novel factors controlling transcriptional memory in *S. cerevisiae*, here we performed a genome-wide screen combining flow cytometry with high-throughput sequencing. We identify multiple genes whose depletion leads to altered gene expression dynamics and focus on the investigation of *RRP6*, a component of the nuclear exosome. The nuclear exosome plays an important role in gene expression and participates in transcription termination^22^, modulates transcription directionality^23,24,25^ and even controls the level of enhancer RNA (eRNAs) in mammals^26^. Like other nuclear RNA degradation pathways, in generally it is thought that susceptibility to degradation by the nuclear exosome is set co-transcriptionally^27^. Although traditionally nuclear decay was thought to affect mainly non-coding transcripts^28^, recent evidence shows that changes in nuclear RNA degradation rates also facilitates gene expression reprograming in response to stress^29,30^. To dissect the potential contribution of the nuclear exosome to transcriptional memory, we characterise gene expression dynamics in wild-type and *rrp6*Δ strains in response to galactose treatment. We categorize genes based on their ability to promote transcriptional memory for both induction and repression of expression in response to galactose. Next, comparing naïve and galactose-primed cells, we investigate the potential contribution of non-coding RNA transcription and chromatin organization to transcriptional memory. We further study the possible direct role of the nuclear exosome modulating mRNA abundance during transcriptional memory. We investigate if differential binding of nuclear surveillance factors could contribute to the faster gene expression reprograming observed in primed cells. Using RNA-crosslinking data, we show that genes with transcriptional memory display a distinct association to nuclear surveillance complexes associated with the nuclear exosome. Finally, we use RNA metabolic labelling to investigate differences in cytoplasmic mRNA turnover between naïve and primed cells. Our results show that changes in nuclear and cytoplasmic mRNA stability between states contributes to differential gene-expression response and transcriptional memory.

## Results

### Genome-wide screening for factors modulating transcriptional memory

To identify genes able to modulate transcriptional memory, we constructed a reporter system able to track this process. As a model for transcriptional memory, we used the expression of *GAL1* in response to galactose in *S. cerevisiae*. Our memory reporter system contains a fast-folding GFP (sfGFP) under the control of the *GAL1* promoter (*pGAL1*), which has been shown to independently display transcriptional memory^4^, and a constitutively expressed MCherry under the control of pTEF1 (see methods and Fig. S1A). Since protein accumulation is delayed in respect to mRNA accumulation, and proteins are in general more stable, we modified both fluorescent reporters with degron signals to facilitate the identification of dynamic changes. The addition of the degron signals increased the resolution of our system by decreasing protein stability and thus minimizing the lag time between mRNA and protein accumulation. We first confirmed the ability of this system to measure transcriptional memory in response to galactose (Fig. S1B). Both raffinose (non-repressed) and glucose (repressed) media have been used as starting point to investigate transcriptional memory in response to galactose^3,45^. Here we decided to use glucose media (YPD), as this simplifies the genome-wide comparison of gene expression between naïve and primed cells (see below).

Next, we transformed the barcoded *S. cerevisiae* deletion collection^31^ with our reporter system, isolated cells based on their *pGAL1-sfGFP* expression using flow cytometry and identified each strain using high-throughput sequencing (see methods for details). To control for technical variability, we included 8 wild-type control strains containing strain-specific barcodes. We grew the pooled barcoded collection in rich glucose media (YPD) and transferred exponentially growing cells (OD_600_ 0.4-0.6, t_0_) to rich galactose media (YPGal) for 3 hours (t_60_, t_120_ and t_180_). After this we transferred cells back to YPD and collected cells after 1.5 (t_90GLU_) and 3 hours (t_180GLU_, which served also as time 0 prime, t_0_’). Finally, we re-exposed cells to galactose for 3 hours and collected a sample each hour (t_60_’, t_120_’ and t_180_’). We sorted cells according to their relative sfGFP expression and identified each strain by sequencing their unique barcode. After correcting for the total number of cells, we generated a virtual *GAL1* gene expression profile over time for each strain (Fig. 1A). By comparing the accumulation of sfGFP between naïve and primed cells we identified potential modulators of *GAL1* transcriptional memory (Supplementary Data 1, see methods for details).

**Fig. 1:**
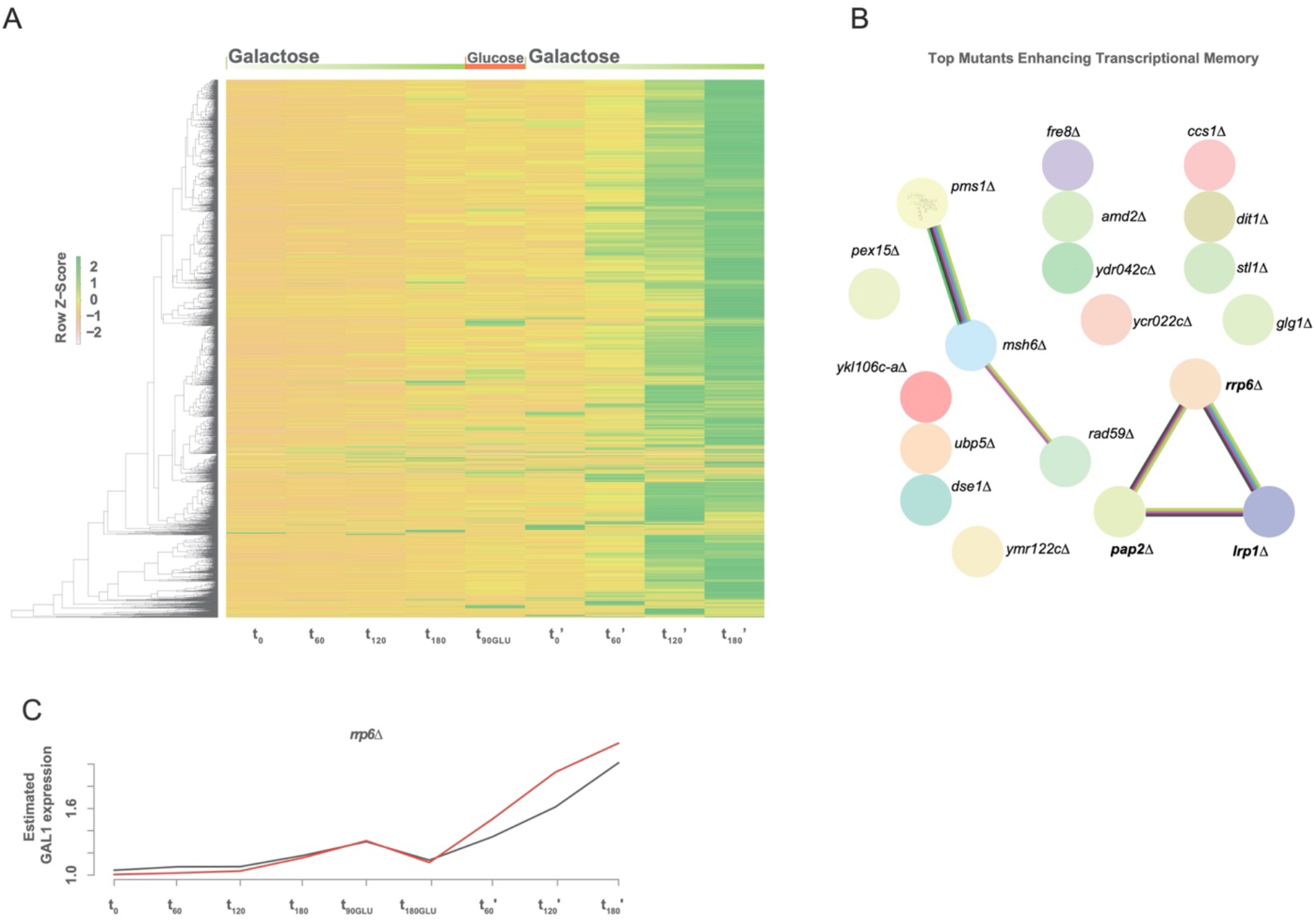
Screen for genetic factors controlling transcriptional memory. **A**, Heatmap depicting strain-specific z-cores for *pGAL1-sGFP* expression at different time points. Green refers to high sfGFP expression (see Supplementary Data 1 and methods for detail). **B**, Top 19 candidate deletion strains displaying enhanced transcriptional memory. Lines represent known interactions as curated by STRING v11.5^40^.blue indicate from curated databases, pink indicate experimentally determined, green indicate predicted interaction as gene neighborhood, light green indicate source as text mining, black indicate co-expression. **C**, Relative sfGFP expression for components of the nuclear exosome (*rrp6*Δ) in red relative to wildtype in black (y-axis from 1 (all cells with no detectable GFP) to 4 (all cells with maximum GFP), see Methods for detail).

We identified 35 mutants with decreased transcriptional memory. Those included mutants for the expected chromatin remodeller ISW2^7^, as well as components of the THO and TREX complexes (*MFT1* and *THP2*) and mitochondrial factors (*ATP18, ATP19, FMP30*…) among others (Fig. S1C). Interestingly, we also identified 37 mutants in which transcriptional memory is enhanced. These included genes involved in nuclear RNA degradation (*RRP6, LRP1* and *PAP2*) and genes involved in meiosis and cell division (*PMS1, MSH6, DSE1*…) (Fig.1B, S1D). Our screen also identified that the depletion of *ELP4* (a member of the elongator complex) enhanced transcriptional memory (Fig. S1D), in agreement with what has been recently described^32^. We decided to study the role of RNA degradation depletion in enhancing transcriptional memory, as it was the unique significantly enriched KEGG term (FDR < 0.0432, using default STRING enrichment). We focused on the role of nuclear mRNA surveillance, since two components of the nuclear exosome (*RRP6* and *LRP1*) were identified in our screen (Fig. 1C). Although the nuclear exosome is well-known to control the abundance of Cryptic Unstable Transcripts (CUTs) in yeast^23,24^, more recent evidence shows that changes in nuclear degradation rates help to rapidly reprogram gene expression after glucose deprivation^29^ and that nuclear decay can dampen the accumulation of full-length mRNAs from stress responsive genes^30^. As the potential role of nuclear mRNA degradation has not been explored in the context of transcriptional memory, we focused on this new avenue.

### Differential transcriptional memory response in absence of functional nuclear exosome

After identifying a potential role of nuclear RNA decay in transcriptional memory using an ectopic reporter system based on protein expression, we validated this result by measuring mRNA expression from the native *GAL1* locus. First, we determined the optimal sampling points to capture the induction/re-induction kinetics at mRNA level by RT-qPCR (Fig. S2A). As expected, optimal RNA sampling points were advanced in comparison to the ones used in our protein based screen (Fig. S2A). As the nuclear exosome has a widespread effect on gene expression^29^, we hypothesized that *RRP6* could modulate transcriptional memory also in genes other than *GAL1*. We performed RNA-Seq in both the wildtype (BY4741) and the *rrp6*Δ strain. We measured mRNA abundance during exponential growth in YPD (t_0_, naïve) and 30 min, 1 hour and 3 hours after change to galactose (t_30_, t_60_ and t_180_). Then we transferred cells to YPD for 3 hours (t_0_’, primed) and measured galactose reinduction at 15 min, 30 min and 1 hour (t_15_’, t_30_’ and t_60_’) (Fig. 2A and Supplementary Data 2). Naïve and primed cell response to galactose was very similar, but primed cells were clearly faster in adapting their transcriptome to the use of galactose as carbon source (Fig. 2B, S2B) in agreement with the existence of transcriptional memory. Specifically, t_0_ and t_0_’ cluster together, while gene expression at t_15_’, t_30_’ and t_60_’ in primed cells clustered with t_30_, t_60_ and t_180_ in naïve cells respectively.

**Fig. 2:**
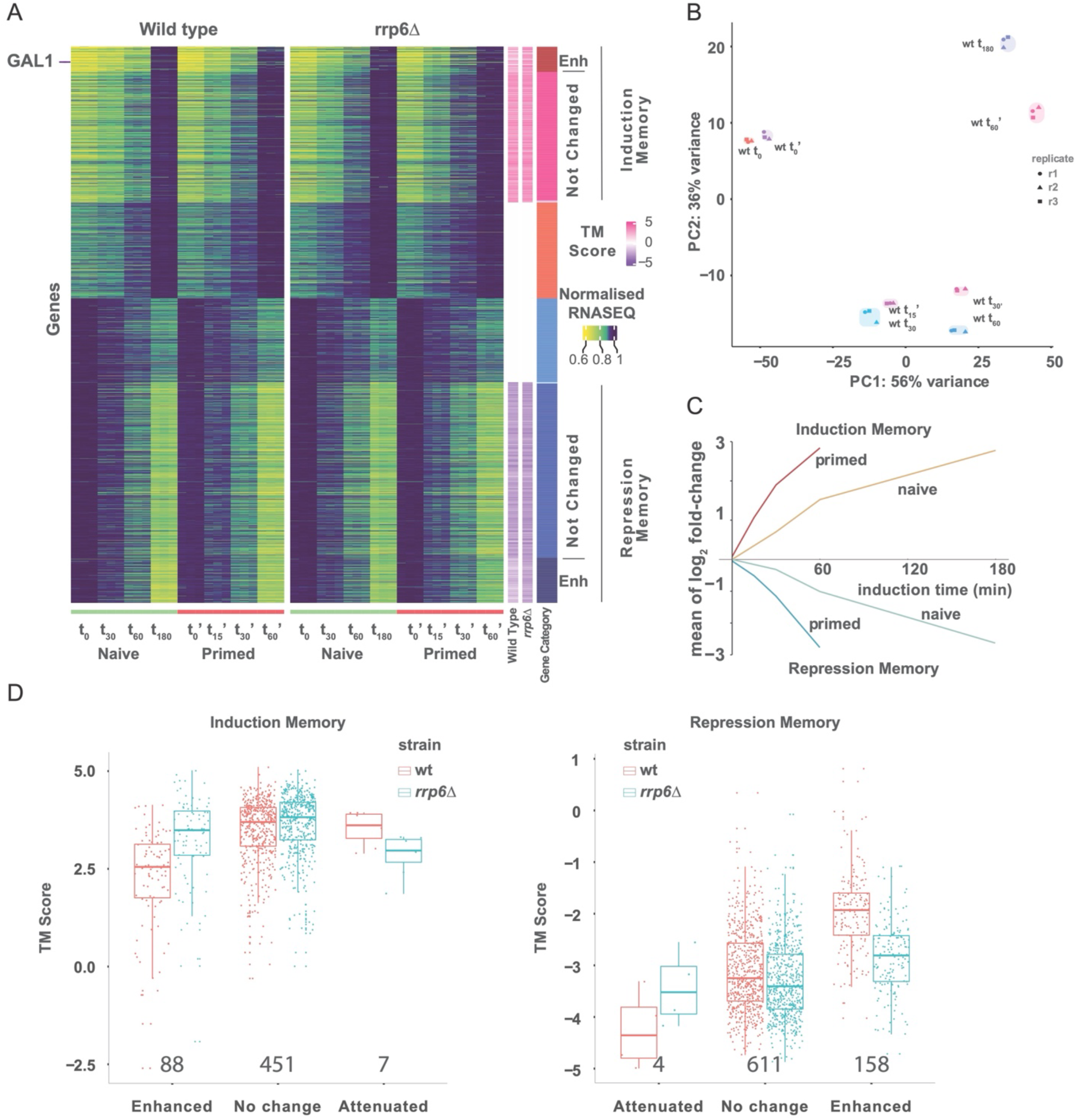
Genome-wide identification of genes with transcriptional memory. **A**, Heatmap of variable genes in RNA-Seq of both wild type and *rrp6*Δ strains. Heatmap intensity represents relative mRNA abundance for each replicate at different timepoints. Gene intensity is normalised by the maximum mRNA abundance across all time points. Gene-specific *transcription memory score* (TM_score_; log_2_ fold-change t_30_’ vs t_180_) is shown in purple to pink for each strain. The rightmost column shows gene category according to TM_score_: induction memory enhanced in *rrp6*Δ (dark red), induction memory unchanged in *rrp6*Δ (red), induced memory attenuated in *rrp6*Δ (pink, very few genes), induced gene with no memory (orange), repressed gene with no memory (light blue), repression memory attenuated in *rrp6*Δ (turquoise, very few genes), repression memory unchanged in *rrp6*Δ (blue), and repression memory enhanced in *rrp6*Δ (dark blue). **B**, Principle component analysis (PCA) of mRNA expression (normalised RNA-Seq data) for both first and second induction in wild type. Only coding mRNAs (ORF-Ts) where considered. **C**, Mean of log_2_ fold-change for genes classified as induction and repression memory genes. Gene expression dynamics in both naïve and primed states are shown. **D**, Memory Score (TM_score_) for genes classified as induction or repression memory.

Focusing on wild-type cells, we identified 882 genes (ORF-Ts, coding transcriptional units as previously defined^24^) induced in response to galactose in both naïve (t_180_) and primed (t_60_’) cells (fold-change > 0, p-adj < 0.001) (Supplementary Data 2 and Fig. S2C). Induced genes contained genes involved in galactose response but also genes related to oxidative phosphorylation and aerobic respiration (Fig. S2D). We also identified 1067 genes which were repressed in response to galactose in both naïve and primed cells (fold-change < 0, p-adj < 0.001) (Supplementary Data 2 and Fig. S2C). Repressed genes include genes associated to ribosome biogenesis, translation initiation and elongation (Fig. S2E). Next, we analysed differential expression along the time course (Fig. S2C and Supplementary Data 2). We quantified transcriptional memory by comparing gene expression dynamics in primed versus naïve cells. Transcriptional memory is commonly used to refer to genes undergoing transcriptional induction, however transcriptional memory also impacts gene repression^33^. Thus, we refer to genes as possessing either induction or repression memory (instead of using the general term of transcriptional memory). We classified genes with induction memory as those with higher induction in primed cells compared to naïve at either 30 minutes or 1 hour (*i*.*e*., induction fold-change in primed state more than 1.5x times the fold-change measured in naïve state). We identified 546 genes with induction memory, including *GAL1* as well as other genes related to ATP metabolic process (Fig. S2F and Supplementary Data 2). On the other hand, we identified 773 genes with repression memory (*i*.*e*., repression fold-change in primed state more than 1.5x times the fold-change measured in naïve state). Repression memory genes were clearly enriched for rRNA processing and ribosome assembly (Fig. S2G and Supplementary Data 2). Despite gene expression between naïve (t_0_) and primed (t_0_’) conditions being similar (Fig. S2C), using external spike-ins we determined that primed cells have a higher proportion of mRNA contribution to the total RNA amount, driven mainly by rRNA abundance (Fig. S2H). This suggests that after 3 hours in glucose, primed cells have adapted their relative mRNA levels to growth in YPD, while rRNA abundance (which has a slower RNA turnover and thus requires more time to reach equilibrium) lags behind.

After investigating mRNA abundance changes in the wild-type strain, we performed the same analysis in *rrp6*Δ. Gene expression changes upon galactose addition in the *rrp6*Δ strain were very similar to the ones observed in a wild-type strain (Fig. S2B). We identified 1026 genes (ORF-Ts) induced in response to galactose in both naïve (at 180 min) and primed (at 60 min) *rrp6*Δ cells (fold-change > 0, p-adj < 0.001), and 1177 genes repressed in response to galactose in both naïve and primed cells (fold-change < 0, p-adj < 0.001) (Supplementary Data 2A). While the main difference between strains relates to the accumulation of CUTs in *rrp6*Δ (Fig. S2I, S2J)^23,24^, we also identified 540 genes differentially expressed between wild-type and *rrp6*Δ strains in the naïve state (p-adj < 0.001). Independently of any difference between wild-type and *rrp6*Δ strains in naïve state, both strains can present transcriptional memory in response to galactose (Fig. 2B, S2B).

Next, we focus on the potential ability of the nuclear exosome to modulate the observed transcriptional memory phenotype. To decrease strain-specific biases, we defined a *“transcription memory score”* (TM_score_) for each strain and gene (Fig. 2D). Specifically, we compared the fold-change after 15 minutes after galactose addition in primed cells (t_30_’) to 3h after galactose addition, t_180_ in naïve cells (which is the maximum fold-change measured for most induced genes in naïve cells). Comparing the transcriptional memory scores between wild-type and *rrp6*Δ strains, we identified 88 genes with enhanced activation memory in *rrp6*Δ (TM_score_ *rrp6Δ /* TM_score_ wild-type > 1.5). As expected from our initial protein-based screen, we confirmed that *GAL1* displayed enhanced induction memory in *rrp6*Δ also in mRNA expression from its native locus. Induction memory genes enhanced by *RRP6* depletion also include other genes involved in aerobic respiration (Fig. S2F and Supplementary Data 2). We identified 451 genes were *rrp6*Δ did not clearly affect the activation memory and 7 genes with attenuated activation memory in *rrp6*Δ (TM_score_ *rrp6Δ /* TM_score_ wild-type < 0.667). Next, we investigated if depletion of the nuclear exosome affected the dynamics of repression memory genes. We identified 158 genes with enhanced repression memory in *rrp6*Δ (TM_score_ *rrp6Δ /* TM_score_ wild-type < 0.667) including ribosomal protein and ribosome biogenesis genes (Fig. S2G and Supplementary Data 2). We identified 611 genes where repression memory was not clearly impacted by RRP6 depletion and only 4 genes with attenuated repression memory (TM_score_ *rrp6Δ /* TM_score_ wild-type > 1.5). Taken together, this shows that depletion of the nuclear exosome can enhance the transcriptional memory both for induced genes (making them increase their mRNA relative abundance faster) and for repressed genes (making them decrease their relative abundance faster).

### Non-coding RNA accumulation or chromatin reorganization do not explain transcriptional memory differences

After showing that multiple genes display an altered transcriptional memory in *rrp6*Δ, we investigated the potential mechanism by which the nuclear exosome could alter this process. Since accumulation of non-coding RNAs (ncRNAs) has been previously associated with a faster reactivation of *GAL* genes from repressive (glucose) conditions^1934^, we investigated if the observed differences in transcriptional memory could be explained by differential ncRNA accumulation. We focused on cryptic unstable transcripts (CUTs) that are well known targets of the nuclear exosome and often arise bidirectionally from coding gene promoters ^2324^. First, we explored if the promoters of genes whose transcriptional memory was enhanced or attenuated after RRP6 depletion overlapped with previously annotated CUTs (*i*.*e*. – 150nt to -50 in the sense strand and -150 to +50 in antisense)^24^. However, we did not observe a clear association between CUTs overlapping promoters and transcriptional memory modulation by the nuclear exosome (Fig. S3A). Focusing on the *GAL1* region, we confirmed the expected gene expression remodelling in response to galactose treatment (Fig. S3B). But we did not observe any clear accumulation of promoter-proximal ncRNA when comparing naïve and primed states. To investigate the potential interaction between non-coding RNA accumulation and transcriptional memory beyond the GAL genes, we investigated relative abundance of non-coding RNAs in naïve and primed states (Fig. S3C). However, we did not observe any clear global difference. Next, we used our transcriptomic data to explore potential differences in unannotated ncRNA transcription overlapping coding genes’ promoters (sense and antisense) depending on their transcriptional memory classification (Fig. 3A). Genes with induction memory enhanced after *RRP6* depletion displayed a clearly increased RNA-Seq coverage in the promoter region in the *rrp6*Δ strain in the sense strand in relation to the wild-type strain. However, no clear differences were observed between naïve and primed states. When investigating antisense transcripts covering promoter regions, we observed a general increase in RNA-Seq coverage for most groups in the *rrp6*Δ strain, as expected from antisense CUT accumulation^23,24^. However, again no clear differences were observed between naïve and primed states. Thus, our results suggest that promoter-proximal ncRNA accumulation is not the main driver in the formation of transcriptional memory.

**Fig. 3:**
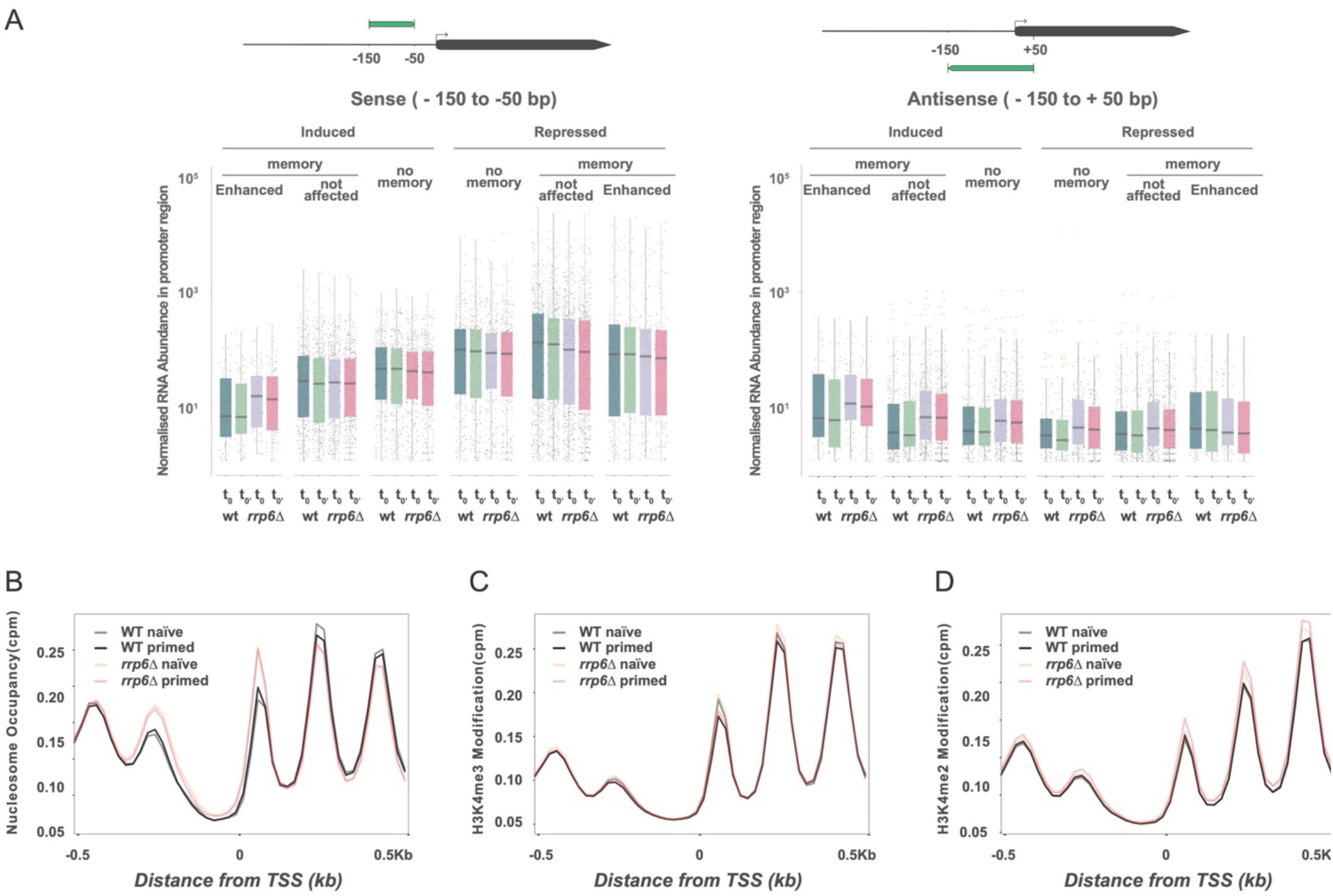
Non-coding transcription and chromatin organization do not explain differences between naïve and primed states. **A**, RNA-Seq coverage of unannotated ncRNA transcription surrounding coding genes’ transcription start sites (TSS) (i.e., overlapping the promoter region). Average RNA coverage at promoter region on sense strand (−150 to -50 bp in respect to TSS) is shown on the left and antisense transcription (−150 to +50 bp in respect to TSS) is shown on the right. Boxplots for wild-type t_0_ naïve (cyan), wild-type t_0_’ primed (green), *rrp6*Δ t_0_ naïve (purple) and *rrp6*Δ t_0_’ primed (pink) are shown. mRNA abundance is normalized to global ORF-Ts abundance. **B**, Global MNase Seq analysis. Metaplot of the distribution of average nucleosome MNase signal. Average sequencing coverage is shown (cpm, counts per million) for wild-type t_0_ naïve (grey), wild-type t_0_’ primed (black), *rrp6*Δ t_0_ naïve (orange) and *rrp6*Δ t_0_’ primed (pink) here. **C**, Global H3K4me3 ChIPseq analysis. Metaplot of the distribution of average H3K4me3 modification signal. **D**, Global H3K4me2 ChIPseq analysis. Metaplot of the distribution of average H3K4me2 modification signal.

Having found no clear association between ncRNA accumulation and transcriptional memory, we investigated potential differences in chromatin. It is well known that chromatin plays an important role in transcriptional memory^35^, however no detailed genome-wide investigation of chromatin organization comparing naïve and primed cells has been conducted in this system. Thus, we performed high resolution nucleosome mapping combining MNase treatment and chromatin immunoprecipitation in naïve (t_0_) and primed (t_0_’) cells for the wild-type and *rrp6*Δ strains. We did not observe any clear difference for global nucleosome occupancy comparing naïve and primed states (Fig. 3B). We only observed a subtle displacement of nucleosomes -1 and +1 towards the nucleosome depleted region (NDR) in the *rrp6*Δ strain in both naïve and primed cells. This *rrp6*Δ-specific alteration is reminiscent of the chromatin remodelling associated to an increase of antisense non-coding transcription that has been recently reported^36^. Next, we investigated nucleosome occupancy for induced and repressed genes (with or without memory), observing similar profiles (Fig. S3D). As subtle changes in chromatin accessibility in *GAL1* promoter have been reported previously^32^, we investigated also that locus specifically (Fig. S3E). However, we did not identify clear differences between naïve and primed cells.

In addition to potential changes in nucleosome occupancy, we also investigated histone marks previously associated to transcriptional memory. Even though in our experimental settings, we focus on short term transcriptional memory to galactose (*i*.*e*., 3h in glucose after initial galactose priming), we decided to explore potential changes in H3K4me3 and H3K4me2 that have been previously associated with long term transcriptional memory (>12 h) for galactose^17^ and heat shock response in plants^35^. Comparing naïve and primed cells at genome-wide level, we observed minimal differences in H3K4me3 and H3K4me2 (Fig. 3C, D). Although in some cases we identified some subtle differences between groups of genes and between strains (Fig. S3D), we did not identify differences between naïve and primed states. This suggests that chromatin differences alone are not sufficient to explain the differential transcriptional memory observed in the *rrp6*Δ strain.

### Differential association of nuclear exosome co-factors modulates transcriptional memory

Having shown that chromatin-mediated regulation was not sufficient to explain the differences in transcriptional memory between wild-type and *rrp6*Δ strains, we focus on the effects of the nuclear exosome modulating mRNA stability. Although traditionally nuclear decay was thought to affect mainly non-coding transcripts^28^, recent evidence shows that changes in nuclear RNA degradation rates can facilitate remodelling of gene expression in yeast^29,30^. These changes can impact gene expression both in a positive and a negative way^29^. Specifically, in response to glucose deprivation, stress responsive genes can better escape nuclear decay (facilitating their accumulation). In contrast, genes downregulated in response to glucose withdraw are targeted more efficiently by nuclear surveillance factors (facilitating their downregulation). Therefore, we hypothesized that the changes in nuclear decay observed in naïve cells could be further enhanced in primed conditions and contribute to the observed *rrp6*Δ-dependent transcriptional memory differences. To test the hypothesis, we investigated if genes with different transcriptional memory behaviour present differential association to the nuclear surveillance complexes TRAMP (Trf4/5-Air1/ 2-Mtr4-polyadenylation) or NNS (Nrd1-Nab3-Sen1). We used CRAC (crosslinking and cDNA analysis) data obtained in naïve conditions measuring the intrinsic association of specific mRNAs to the TRAMP (*i*.*e*. Mtr4) and NNS (*i*.*e*. Nab3) complexes^29^. We observed that genes induced after galactose addition generally exhibit higher intrinsic association to both complexes, while genes repressed in response to galactose are relatively less bound (Fig. 4A, B). Interestingly, this trend was even more clear when analysing genes with induction and repression memory. Specifically, induced genes with transcriptional memory (for which mRNA abundance increases faster in primed cells) are more associated to TRAMP and NNS complexes, while repressed genes with transcriptional memory are less associated (Fig. 4C, D). Next, we explored up to what degree differential association to the nuclear surveillance complexes could explain the transcriptional memory phenotype observed in the *rrp6*Δ strain. We observed that genes where induction memory was enhanced by the nuclear exosome depletion had in general higher association to TRAMP and NNS (Fig. 4E, F). On the contrary, repressed genes whose repression memory was enhanced by the nuclear exosome depletion had in general lower association to NNS (Fig. 4G, H). Thus, our analysis suggests that differential binding to nuclear surveillance factors, and potentially changes in nuclear RNA degradation rates, contribute to transcription memory.

**Fig. 4:**
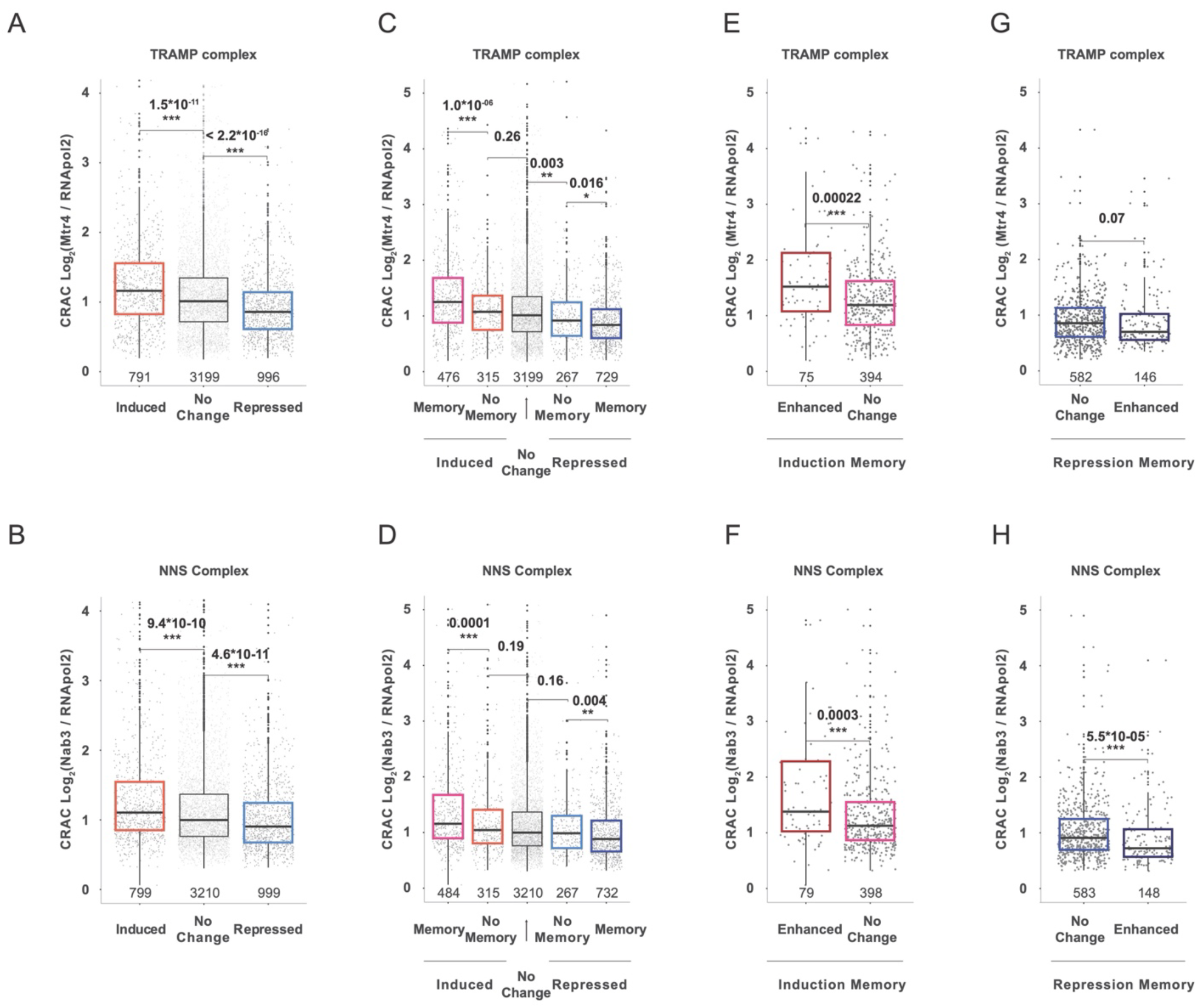
Differential association to nuclear exosome co-factors. **A**, Relative association for the TRAMP complex (i.e. Mtr4) as measured by CRAC in naïve cells ^29^. Boxplot for induced and repressed genes is shown. To measure gene-specific nuclear surveillance association, since those complexes act on nascent transcripts, CRAC data is normalized by RNA pol II association, as previously described. Number of analysed genes is indicated in grey. **B**, Relative association for the NNS complex (Nab3) as measured by CRAC in naïve cells. **C-D**, as in A but for genes classified according to their transcriptional memory. **E-F**, As in A but comparing genes with induction memory enhanced or not enhanced by RRP6 depletion. **G-H**, As in A but comparing genes with repression memory enhanced or not enhanced by RRP6 depletion. Significance computed using Wilcoxon signed-rank test.

### Changes in cytoplasmic RNA stability also modulate transcriptional memory

Having investigated the role of nuclear RNA degradation in transcriptional memory, we decided to study the potential role of other RNA degradation pathways. We reasoned that, as targeting for nuclear RNA degradation is often set co-transcriptionally^27^ changes in nuclear mRNA stability will mainly modulate the appearance of newly synthetised molecules. However, it could be expected that, to modulate the abundance of mRNAs already presents in the cytoplasm (*e*.*g*. genes with repression memory), cytoplasmic mRNA decay would be also altered between naïve and primed cells. Unfortunately, key components of the cytoplasmic mRNA decay such as *XRN1* lacked sufficient coverage to be included in our analysis (Supplementary Data 1). To investigate if global changes in mRNA stability (and not only nuclear decay) contribute to transcriptional memory we measured genome-wide mRNA stability using metabolic RNA labelling (SLAM-Seq)^37,38^ in naïve and primed cells. SLAM-Seq compares total mRNA abundance to that of new mRNA molecules generated during the metabolic RNA labelling pulse. Thus, it can be expected that it captures mainly changes in mRNA stability due to differences in cytoplasmic mRNA decay, while changes in nuclear decay would be difficult to measure (as nuclear RNA decay acts mainly on nascent mRNA molecules). We labelled newly synthetized RNA with thiouracil for 10 minutes and harvested cells at 0 and 30 min after galactose addition for both wild-type and *rrp6*Δ strains (see methods). As expected upon shift to galactose, generation of newly synthesized mRNA molecules was drastically decreased (Fig. S4A). Thus, we focus our analysis on naïve (t_0_) and primed (t_0_’) conditions where we could assume a steady state between mRNA synthesis and decay (*i*.*e*. synthesis rate is in equilibrium with the mRNA decay rate).

Investigating the wild-type strain, we observed a clear decrease in mRNA turnover in the primed condition, indicating a general stabilization of the cellular mRNAs (p-value < 2.2·10^−16^, Figure 5A). Although most mRNAs increase their mRNA stability in primed cells (slower turnover), genes associated to respiration and galactose metabolism were particularly stabilized (Fig. S4B and Supplementary Data 3). On the other hand, genes associated with cytoplasmic translation were less stabilized (Fig. S4C and Supplementary Data 3). Next, we investigated the changes in mRNA turnover between naïve and primed conditions for different groups of genes. Focusing on genes induced after galactose treatment, we observed that their mRNA stability was significantly increased (slower turnover) in primed cells (Fig. 5B), something that could facilitate their accumulation upon re-induction. Stability of induced genes with transcriptional memory was increased even more (Fig. 5C and S4D, p-value < 2.8·10^−6^). On the contrary, the mRNA stability of genes repressed after galactose treatment increased less than the global population (i.e., less decrease in mRNA turn-over, Fig. 5B). And again, the trend was even more clear for mRNAs with repression memory (Fig. 5C and S4D). In agreement with our model, maintaining a relatively faster turnover (lower mRNA stability) in primed cells could contribute to the fast decrease of their mRNA abundance upon transcription shut down. Next, we investigated how the stability of repression memory genes enhanced by *RRP6* depletion, changed between naïve and primed conditions. We observed a decrease in stability in respect to other repression memory genes (Fig. 5D), suggesting that changes in mRNA stability contribute to the observed transcriptional memory phenotype. Genes with induction memory enhanced by *RRP6* depletion presented a subtle relative increase in mRNA stability in primed cells, but this change was not significant (Fig. S4E). This could be caused by a different regulation mechanism or due to our limited ability to accurately measure their mRNA turnover in non-induced conditions (where their mRNA abundance is extremely low).

**Fig. 5:**
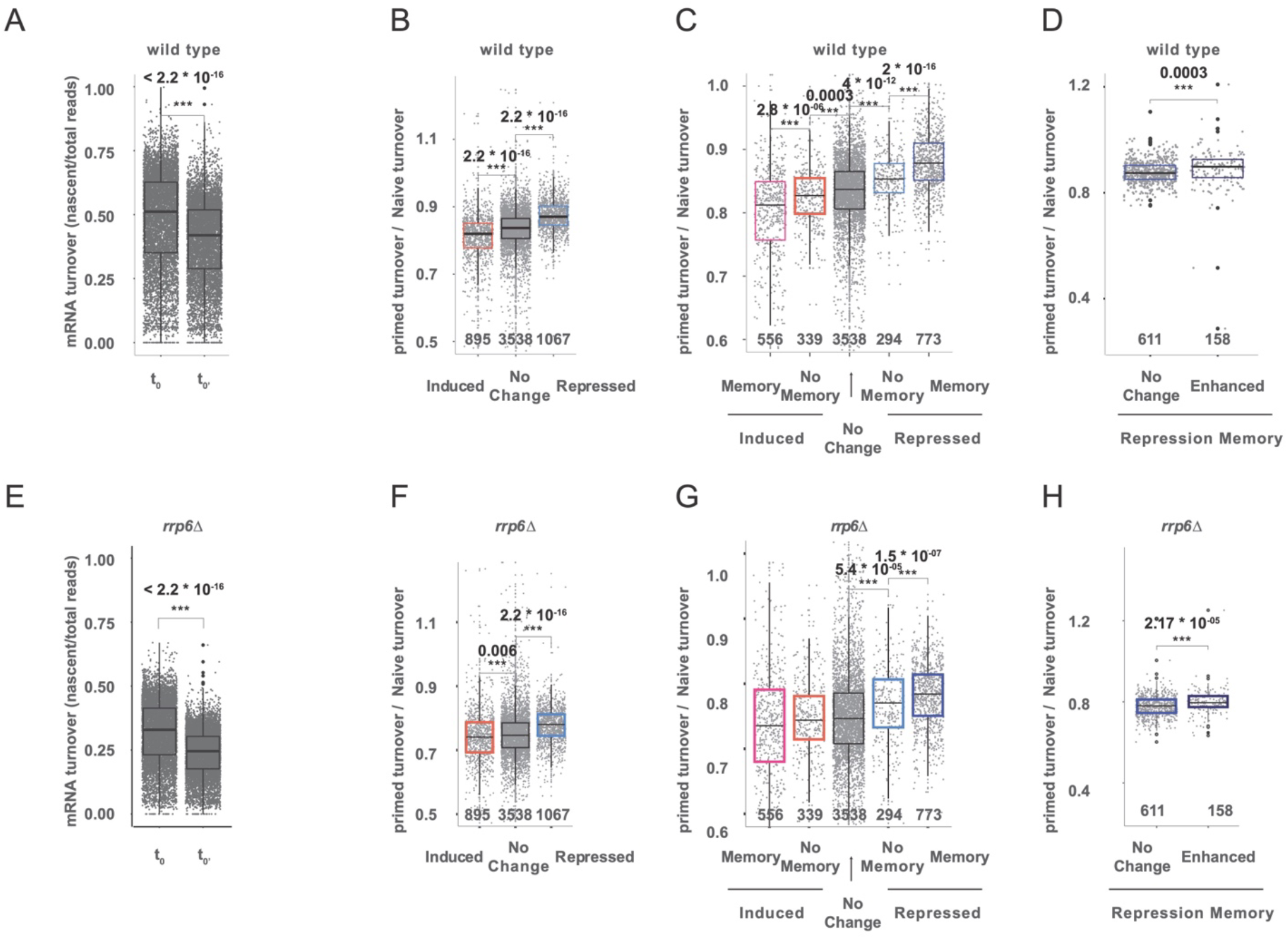
Differential mRNA turnover between naïve and primed cells. **A**, Relative mRNA turnover (comparing nascent vs total RNA) using SLAM-seq in naïve (t_0_) and primed (t_0_’) conditions. High turnover indicates faster transcription/decay and thus lower mRNA stability. Gene-specific turnover was computed by comparing for each gene the reads containing T>C conversion to the total mapped reads. Only reads containing at least 2 T>C conversions are considered newly synthesized reads. Only genes with at least 20 total reads were considered for RNA turnover analysis. **B**, Change in mRNA turnover between primed and naïve conditions for induced and repressed genes in response to galactose. Number of analysed genes is shown at the bottom of each boxplot. **C**, As in B but for genes according to their transcriptional memory profile. **D**, As in B but for genes with repression memory enhanced or not enhanced after RRP6 depletion. **E-H**, As A-D but for the *rrp6*Δ strain.

Finally, we expanded our analysis to investigate global changes in mRNA stability in the *rrp6*Δ strain. Using SLAM-Seq, we determined that *rrp6*Δ has a slower mRNA turnover than the wild-type strain (Fig. 5E, p-value < 2.2·10^−16^). As expected, this effect was particularly strong for CUTs, which were clearly stabilized in *rrp6*Δ (Fig. S4F, p-value < 2.2·10^− 16^). Having validated our RNA turnover measurements in *rrp6*Δ, *we* investigated differences in mRNA turnover between naïve and primed conditions. As in the wild-type strain, we observed a clear decrease in mRNA turnover in primed *rrp6*Δ cells (Fig 5E, S4A, p < 2.2·10^−16^). This shows that the observed mRNA stabilization in primed cells (decreased mRNA turnover) does not depend exclusively on the nuclear exosome, but that regulation of the cytoplasmic mRNA decay is also involved. In agreement with that observation, RNA turnover regulation between naïve and primed conditions was similar to that of the wild-type strain. Specifically, genes displaying induction memory underwent a significant increase in their relative mRNA stability (slower turnover) in primed cells, while genes with repression memory decreased in their relative RNA stability (faster turnover) in primed cells (Fig. 5F-H, S4G-H). To further dissect the contribution of cytoplasmic factors to this process, we investigated if genes with differential transcriptional memory phenotype were altered for this process. Codon optimality has been shown to be a main player controlling cytoplasmic mRNA stability^39^. Interestingly genes with repression memory enhanced by depletion of the nuclear exosome present a particularly high codon adaptation index (Fig. S4I). Taken together, our results show that changes in mRNA stability between naïve and primed states contribute to the transcriptional memory phenotypes and that multiple factors, and not only the nuclear exosome, likely take part in this process.

## Discussion

Previous studies investigating transcriptional memory have focused on how direct modulation of the transcription process enables faster gene re-induction kinetics. However, even though steady-state mRNA levels reflect the combination of transcription and RNA decay, the role of mRNA degradation in this process remained unexplored. Here we show that differential regulation of mRNA stability facilitates gene expression adaptation in response to environmental changes and that this process can be influenced by previous stimuli.

In this study, we performed a genome-wide screen to identify novel factors able to modulate transcriptional memory in budding yeast in response to galactose as carbon source. In addition to confirming the role of known players in this process, we show that the depletion of the nuclear exosome leads to faster reinduction kinetics of the *GAL1* gene in galactose-primed cells. To dissect this process, we performed a detailed RNA-Seq study of gene expression changes in naïve and primed cells. Consistent with previous work ^33^, we identify that transcriptional memory both facilitates the induction of particular genes (activation memory) and also accelerates the repression of others (repression memory). Unexpectedly, when investigating the role of the nuclear exosome in this process, we discovered that its depletion could enhance both activation and repression memory. Specifically, *rrp6*Δ depletion enhanced the transcriptional memory of a subset of induced genes (making them increase their mRNA abundance faster) and also the repression of a different subset of genes (making them decrease mRNA abundance faster). The groups displaying enhanced transcriptional memory phenotypes were not random. For example, genes related with carbohydrate metabolism or meiosis displayed enhanced activation memory in *rrp6*Δ, while cytoplasmic translation was enriched in the group of genes with enhanced repression memory (Fig. S2 and Supplementary Data 2). As the nuclear exosome has a well-known role in controlling promoter directionality, transcription termination and the abundance of non-coding transcripts (*i*.*e*., CUTs), we first investigated its potential role in controlling the transcription process. However, our analysis discarded a potential indirect role of the nuclear exosome through controlling the accumulation of CUTs overlapping promoters of coding genes in this process ^5^. We also discarded potential chromatin differences when comparing naïve and primed states.

Next, as our result showed that the depletion of the nuclear exosome led to divergent effects in groups of genes that are regulated in opposite directions during cellular adaptation, we considered if the effect could be mediated by regulation of mRNA decay (instead of changes in the transcription process). Recent work shows that the nuclear exosome, in addition to its canonical role in regulation of ncRNA abundance, is also important in facilitating remodelling of gene expression in yeast^29,30^. With those works as a starting point, we investigated if, in addition to its role in modulating coding mRNA abundance during the stress response, the nuclear exosome could also behave differently in naïve and primed conditions. Our analysis showed that genes whose induction memory was enhanced by the nuclear exosome depletion had in general higher association to nuclear surveillance factors (*i*.*e*., NNS and TRAMP complexes). On the contrary, repressed genes with memory enhanced by the nuclear exosome depletion had in general lower intrinsic association to these factors. This suggested a scenario where the nuclear exosome could limit the accumulation of mRNAs from genes with activation memory, while having almost no effect in mRNAs from genes with repression memory. This would explain our observations, since in absence of the nuclear exosome, mRNAs from genes with activation memory would accumulate faster. On the contrary, nascent mRNAs from genes with repression memory would not be particularly stabilized in absence of the nuclear exosome. However, as most mRNAs would undergo a subtle stabilization in absence of the nuclear exosome, this would mean that mRNAs from genes with repression memory would decrease in their relative abundance faster than the global population.

Finally, since the contribution of nuclear decay can be expected to be relatively small in comparison to cytoplasmic decay, we investigated if global changes in mRNA stability (and not only nuclear decay) could contribute to differences between naïve and primed cells. We reasoned that changes in cytoplasmic mRNA stability would be especially important in facilitating a faster downregulation of mRNAs of genes with repression memory, as in those genes transcriptional activity, and consequently nascent nuclear mRNA decay, can be expected to be low. To test this hypothesis, we measured RNA stability in primed and naïve cells using RNA metabolic labelling. Despite naïve and primed cells displaying an almost identical gene expression program (Fig. 2, S2), our results clearly show that primed cells undergo a decrease in mRNA turnover, indicating a general stabilization of the cellular mRNAs (Fig. 5A). However, this stabilization is not equal across all gene groups. In fact, genes that are induced after galactose treatment exhibit a significantly higher mRNA stability in primed cells, something that would facilitate their accumulation upon re-induction. This stabilization was even more pronounced for genes with activation memory. On the contrary, genes that are repressed after galactose treatment exhibit a lower relative increase in their mRNA stability in primed cells in respect to the global population. This relative destabilization was even more clear for genes with repression memory, something that would facilitate their faster decrease upon galactose re-stimulation. Interestingly, although our work shows involvement of *RRP6* in regulation of stability in a subset of repression memory genes, we also observe clear changes in mRNA stability between primed and naïve cells even in the absence of the nuclear exosome. Interestingly those genes with higher relative destabilization In primed conditions present a particularly high codon adaptation index (Fig. S4I). This suggest that they could be more sensitive to drastic changes in translation (and thus co-translationally regulated cytoplasmic mRNA stability). However, more research would be required to investigate this possibility.

Taking all this together, our work suggest that there is a co-regulation between the transcriptional responses associated to faster transcriptional induction and repression and the associated changes in post-transcriptional mRNA life (at nuclear and cytoplasmic levels). This suggests a general connection between transcription rate and nuclear and cytoplasmic mRNA decay, which together modulate mRNA abundance. Thus, our work suggests the existence of a coordination between “*transcriptional*” and “*post-transcriptional memory*” to facilitate swifter adaptation to changing environments. Although here we have focused on the dissection of gene expression during carbon-source change in budding yeast genes, we anticipate that similar synergistic transcription-mRNA stability crosstalk could occur in other conditions where massive gene expression changes are expected. Understanding how transcription activity and mRNA decay can change while maintaining apparently identical mRNA abundance will be important to understand differential behaviour in adaptation to changing environments and cellular response.

## Acknowledgements

We thank all members of the Pelechano, Kutter and Friedländer laboratories for useful discussions. We thank Adrian Cortés-Sanchón for initial support with the transcriptional memory screening and Yujie Zhang for technical assistance. We thank José Enrique Pérez-Ortín, Yerma Pareja and Bastian Linder for useful comments in the manuscript. Computational analysis was performed on resources provided by the Swedish National Infrastructure for Computing (SNIC) through Uppsala Multidisciplinary Center for Advanced Computational Science (UPPMAX) partially funded by the Swedish Research Council through grant agreement no. 2018-05973. We also acknowledge the use of the HPC Cloud Platform of Shandong University.

## Author Contributions

VP and BL conceived the study. PZ, MMT and GL performed and analyzed the screen with supervision from VP and LMS. AA contributed to the chromatin analysis. BL performed all other experimental and computational work. BL and VP drafted the original manuscript. All authors reviewed and edited the manuscript. VP supervised the project.

## Methods

### Generation of a genome-wide reported for pGAL1 transcriptional memory

To generate the pGAL1 memory reporter system we used p416 TEF as backbone^41^. The final plasmid contains a constitutively reporter with a pTEF promoter expressing mCherry-degron and a CYC1 terminator. The galactose reporter is controlled by a pGAL1 promoter expressing sfGFP-degron and an ADH1 terminator. The plasmid confers Nourseothricin resistance (NAT1) to enable its transformation into the barcoded yeast deletion collection containing KanMX^31^ (Fig S1A). To obtain repeated measure of a wild-type strain in response to glucose-galactose transition we generated 8 barcoded control strains where we introduced a KanMX cassette containing strain-specific barcode next to the ura3Δ0 locus in a BY4741 strain. We transformed the diploid deletion^31^ collection with the pGAL1 memory reporter plasmid using Gietz’s Frozen competent yeast transformation protocol^42^. The pooled collection was grown overnight in YPD (+200µg/ml Geneticin) and processed in exponential phase OD_600_ ∼ 0.5. To increase sample complexity and minimize cell bottlenecks we performed multiple transformations. For each pool transformation we used 10^8^ cells and 236 µg plasmid DNA. The heat shock was performed for 1h at 42°C in a heat-block with 5 min 1000rpm interval shacking and additional vertexing every 20 min. Antibiotic selection was performed on Nourseothricin(60µg/ml) plates to minimize competition between adjacent clones. We performed a total of 15 independent transformations that led to ∼30,000 clonal colonies. Colonies were scratched, mixed with glycerol (final 21%), aliquoted and frozen. For the 8 barcoded control strains we performed the same process but transforming independently each strain.

### Reporter based screen for transcriptional memory

Frozen aliquots of the barcoded genome-wide deletion collection barcoded controls containing the reporter system were revived for 8.5-10 hours in YPD+ Nourseothricin (60µg/ml). We performed in parallel the experiment with the pooled genome-wide deletion collection and the pooled barcoded controls. When cells reached exponential growth (OD_600_ 0.4-0.6), we took and aliquot and defined that point as TP0 (naïve). We changed media to galactose (YPGal) for 3 hours and collected an aliquot every hour (TP1= t_60_, TP2= t_120_, TP3= t_180_). When then changed cells back to glucose media for 3 hours and collected TP4 (t_90GLU_) and TP5 (t_180GLU_ which served also as t_0_’). Finally, we re-exposed cells to galactose for 3 hours and collected a sample each hour (TP6= t_60_’, TP7= t_120_’, TP8= t_180_’). Media change was performed by spinning cells (5 min 2500g) and performing one wash with the new media before final resuspension. For fluorescent detection cells were fixed in 50% ethanol (final) and stored at 4°C. Fixed pooled barcoded controls and pooled genome-wide deletion were combined at 1:65.5 cell ratio (every barcoded control would correspond to 1:500 of all cells). Before fluorescent measurement, cells were spun down (5 minutes 3000g), washed with PBS and spun down for 30 minutes to allow the sfGFP to refold. After this cell were resuspended in PBS for FACS.

For FACS sorting we fixed gates to consider only those cells significantly expressing MCherry. After this filter, we split sfGFP expression in 4 gates: no expression (R5), low expression (R6), intermediate expression (R7) and high level of sfGFP expression (R8). We aimed to sort a minimum of 10^6^ cells per gate when feasible, and sorted between 7 and 25 million cells per time point (see DataS2 for details). We performed two independent replicates.

For DNA isolation cells were resuspend in 300 μL of 10’prep buffer (2% TritonX-100, 1% SDS, 100 mM NaCl, 1 mM EDTA y 10 mM Tris-HCl pH 8) and transferred to a 2 mL tube with 500 μL Phenol:Chlorophorm:isoamilic (25:24:1) and 500 μL glass beads. Cells were vortexed using 4 30 second pulses in a FastPrep-24 (MP Biomedical) instrument at 5.5m/s. We recovered the aqueous phase after centrifugation and a gel-lock tube with 500 μL Phenol:Chlorophorm:isoamilic (25:24:1). We vortexed the sample, centrifuge and recovered the aqueous phase. We added 3 L RNase A (DNase-free) (10 mg/mL) and incubate during 30 min at 37°C. Ilumina compatible sequencing libraries of the UpTag region of each strain were generated by two consecutives PCRs. During the first PCR we used oligos indexing for the biological replicate (KanMX) and the time points (UPTAG). During the second PCR we introduced illumine compatible oligos and indexed for sGFP signal (see DataS2 for details). We performed 4 independent PCRs for each sample. In brief, for the first PCR reaction contained 2µL extracted gDNA, 1 µl dNTPs (10 mM), 0.25 µl UpKanMX primers (10 mM), 0.25 µl UPTAG primers (10 mM), 10 µl 5 x Phire Reaction Buffer, 0.3 µl Phire Hot Start II DNA Polymerase (Thermo Fisher Scientific) in a total reaction volume of 50 µl. PCR program was conducted for 30 s at 98 °C, 15-16 cycles of 10 s at 98 °C, 10 s at 63 °C, 30s at 72 °C and final elongation for 5 min at 72 °C. For the second PCR we used 2µL product from the first PCR, 1 µl dNTPs (10 mM), 0.5 µl PE2_MPX (10 mM), 0.5 µl PE1.0 (10 mM), 10 µl 5 x Phire Reaction Buffer, 0.3 µl Phire Hot Start II DNA Polymerase (Thermo Fisher Scientific) in a total reaction volume of 50 µl. PCR program was conducted for 30 s at 98 °C, 15-16 cycles of 10 s at 98 °C, 10 s at 65 °C, 30s at 72 °C and final elongation for 5 min at 72 °C. PCR replicates were pooled, purified with HighPrep beads (MagBio), quantified and pooled at equal concentration. Libraries were sequenced using the Illumina HiSeq2000 platform.

### Transcriptional memory screen analysis

To assign mapping sequenced barcodes onto strains we constructed a fasta contig for each deletion strain with its uptag and downtag, and these fasta contigs were indexed using bwa. The sequencing reads were trimmed down to the first 45 bp and then aligned using bwa aln (default settings with –B 6) and bwa se (default settings with –n 1). The first 6 bp of the barcode that indicates the timepoint of the sample was placed under the BC tag during alignment with bwa (the –B 6 option). After alignments, counts for each deletion strain in each timepoint and window were tabulated using R.

We focus on the comparison of the differences between GFP accumulation in naïve (TP3-4) and primed (TP5-6) cells. Please note that, as protein accumulation is delayed respect to mRNA production, TP4 correspond to cells already transitioned back to glucose.

To test for differences in memory response we modelled the change in barcode counts for each strain using a quasibinomial generalized linear model. For each transition (e.g TP3 to TP4), for timepoint i, biological replicate j and strain k

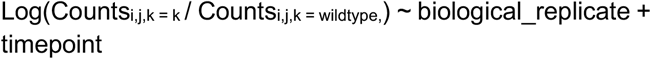

The extra dispersion parameter for quasibinomial glm is estimated from the data using Pearson’s coefficient of dispersion (implement by quasibinomial() in R)

For transition TP3 to TP4, we can obtain an estimate of the change in log odds ratio between TP4 and TP3, β_1_ and also an estimate of its error, se_1_. Likewise in transition from TP5 to TP6, we obtain an estimate β_2_ and se_2_. To ask if the responses during the second transition (TP5 to TP6) is significantly different from the first transition (TP3 to TP4), we calculate a z-score for this:

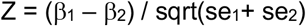

Then we combined the z scores for 3 windows using Stouffer’s method, using the cell counts in each windows as weights. It is to be noted that we take the absolute of all Z values since they can take different signs. After calculating this estimate of Stouffer Z, we check whether the z scores change signs more than once before calling it a significant change in responses. We identify 35 mutants with putative decreased transcriptional memory (TP5_6 < TP3_4) and 37 with enhanced transcriptional memory (TP5_6 > TP3_4) (Supplementary Data 1)

To calculate GFP scores of individual strains at the different timepoints, several data manipulation were performed. First, a cutoff was applied to only include abundant strains (rowMeans > 5). To obtain the relative abundance of the strain in the corresponding sample, for each sample, reads of individual strains were normalized by dividing them with the total number of their samples. Wild type and deletion strains have shown a dispersed distribution across the 4 GFP windows and across time points. In order to obtain the averaged GFP expression of individual strains at the different time points, the contribution of each of 4 GFP windows to average GFP expression has to be normalized. Thus, relative numbers of FACS sorting events for the 3(4) different GFP fractions per time point were multiplied with the corresponding relative reads of individual strains at the different time points. Next for each GFP window a score was assigned (GFP^-^=1, GFP^low^=2, GFP^mid^=3, GFP^high^=4). Normalized reads were multiplied with their respective GFP score, summed up for individual strains at individual time points and divided by sum of the non-multiplied values.

### RNA-Seq experiment and library preparation

*Saccharomyces cerevisiae* strain BY4741 (MAT a *his3Δ1 leu2Δ0 met15Δ0 ura3Δ0*) was grown to exponential phase (OD600∼0.5) in YPD medium (1% yeast extract, 2% peptone, 2% glucose) for at least 16h at 30°C (naïve cells). To change cells from YPD to YPGal (1% yeast extract, 2% peptone, 2% galactose) cells were collected by 2 min centrifugation (3000g) and washed with prewarmed YPGal. After wash, the cells were collected by 2 min centrifugation (3000g) and resuspended in prewarmed YPGal for 3 hours. Next cells were shifted to glucose containing media (YPD) for 3 hours, performing a wash with prewarmed YPD as previously described. Finally, galactose primed cells where washed and exposed to prewarmed YPGal. All yeast samples (2ml) were collected by centrifugation (30 seconds at 8000 x g) and pellets were frozen in liquid nitrogen. *Schizosaccharomyces pombe* (h-) used as spike-in was grown at 30°C to mid-log phase (OD600 ∼0.5)

For library construction, total RNA concentration was measured with Qubit and quality by capillary electrophoresis. We used 2.5 μg total *S. cerevisiae* RNA supplemented with 0.6ng SIRV-SET3 (Lexogen) as spike-in. rRNA was depleted with illumina Ribo-Zero Gold rRNA Removal Kit (Yeast) according to manufacturer instructions. Then, a strand-specific RNA-Seq library was prepared using NEBNext Ultra Directional RNA Library Prep Kit for Illumina following the manufacturer instruction. Briefly, rRNA depleted RNA was first fragmented and then we use random primer to generate first cDNA strand. dUTP was incorporated into cDNA during the following second strand synthesis. After end repair and dA tailing, Illumina adaptors are ligated. The second strand containing dUTP was removed using USER enzyme mix. Strand specific library was prepared with 7 PCR cycles. Library quality was assessed via Qubit and Bioanalyzer znd sequenced using an Illumina Nextseq 500 instrument.

### Processing, analysis and graphic display of RNA-seq data

The genome assembly and annotation for the RNA-Seq data analysis was downloaded from SGD database (version 64-1-1) and annotation from (Xu et al., 2009). The quality of the RNA-Seq data was assessed with FastQC (Andrews, 2010). Reads were aligned to the transcriptome by STAR with parameters “--outFilterMismatchNmax 4 -- alignIntronMin 13 --alignIntronMax 2482”.

Reverse stranded reads were then summarized into gene expression values by featurecounts with parameter “-s 2 - C”. Chimeric fragments were excluded from fragment counting.

Read counts were normalized by sum of coding transcriptome. Differential gene expression analysis was perfomed using the DESEQ2 package in R. When calculating log fold change of different time points, we use the time 0 of first induction as reference point.

PCA plot data was calculated with plotPCA function of DESEQ2 package then plotted with ggplot2. MA plot data was calculated with plotMA function of DESEQ2 and then plotted with ggplot2.

Stepwise Annotation for the gene category: first sort gene into 3 groups(Induced genes, genes with no change and repressed genes) then into 5 groups according to memory pattern (induction memory genes, induced genes without memory, genes with no change, repressed genes without memory and genes with repression memory).

Induced genes are defined as genes whose (lfc, log_2_ fold change) lfc^3h^ > 0, lfc^1h^’ > 0, lfc^3h^ > lfc^1h^, lfc^3h^ > lfc^30min^ and adjusted p value < 0.001. Induction memory genes are defined as induced genes whose lfc^30min’^ - lfc^30min^ > log_2_(1.5) or lfc^1h^’ – lfc^1h^ > log_2_(1.5). Repressed genes are defined as genes whose lfc^3h^ < 0, lfc^1h^’ < 0, lfc^3h^ < lfc^1h^, lfc^3h^ < lfc^30min^ and adjusted p value < 0.001. Repression memory genes are defined as those repressed genes whose lfc^30min’^ - lfc^30min^ < -log2(1.5) or lfc^1h^’ – lfc^1h^ < -log2(1.5). lfc^1h^’ and lfc^30min’^ are the log 2 fold change of 1h and 30 min in second induction (primed state) while lfc^30min^, lfc^1h^ and lfc^3h^ are the log fold change of 30 min, 1 hour and 3 hours in the first induction (naïve state). For memory genes affected by *RRP6* depletion, we first normalized the gene log fold change in primed state by the gene log fold change of longest induction time point (3h) in naïve state to cancel the effect of mutation and have fair comparation of memory effect between strains.

Heatmap was generated with Complexheatmap. TM_score_ was defined and calculated as a measure of relative change amplitude at 15 min in primed state normalized by RNA abundance of 3h in naïve state. For induced memory genes, memory index is illustrated as lfc^15min^’ – lfc^3h^ + 5. For repression memory genes, memory index is illustrated as lfc^15min^’ – lfc^3h^ – 5.

Bigwig file for IGV visualization was generated with deeptools bamcoverage with normalization factor generated by inverting size factor generated in DESEQ2 normalised by sum of coding transcriptome. Hypergeometric test was performed using https://systems.crump.ucla.edu/hypergeometric/. GO enrichment was performed with R package “clusterProfiler”. mRNA codon stability index, translation efficiency were obtained from Carneiro et al^43^.

### MnaseSeq and ChIPseq experiment and library preparation

We analyzed *S. cerevisiae* cells (wildtype and *rrp6*Δ) and used *S. pombe* as spike-in. Cell cultures were grown until OD_600_ 0.3-0.5 (*S. cerevisiae*) or 0.74 (*S. pombe*). 100 ml culture per *S. cerevisiae* sample and 30 ml of S. *pombe* culture was crosslinked using 1% formaldehyde for 15 minutes at room temperature. Formaldehyde was quenched by 0.125 M glycine for 5 minutes. Then cells were washed three times with cold TBS, flash-frozen in liquid nitrogen and stored at -80 °C. Frozen cell pellets were resuspended in zymolyase solution (1 M sorbitol, 50 mM Tris-HCl pH 7.5, 1% beta-mercaptoethanol, 0.1 U/µl zymolyase). The zymolyase digestion proceeded at +37 °C for 30 min (*S. cerevisiae*) or 90 min (*S. pombe*). Spheroplasts were isolated by centrifugation at 6000xg for 10 minutes at +4 °C. Spheroplasts were resuspended in NP buffer (10 mM Tris-HCl pH 7.5, 1 M sorbitol, 50 mM NaCl, 5 mM MgCl_2_, 1 mM CaCl_2_, 0.075% NP-40 (Tergitol), 1% beta-mercaptoethanol, 0.5 mM spermidine, 1% yeast protease inhibitor cocktail). The suspensions were pre-warmed and then 0.5 U/µl of MNase for *S. cerevisiae* and 2 U/µl for *S. pombe* was added per sample. MNase digestion proceeded at 37 °C for 40 min (*S. cerevisiae*) or 30 min (*S. pombe*) and stopped by addition of EGTA. The supernatants containing chromatin fragments were collected and diluted with 1 ml RIPA buffer (10 mM Tris-HCl pH 8.0, 1 mM EDTA pH 8.0, 0.1% SDS, 140 mM NaCl, 1% Triton X-100, 0.1% sodium deoxycholate) with 1% yeast protease inhibitor cocktail. Equal volumes of *S. pombe* chromatin were spiked in to *S. cerevisiae* chromatin samples at this point, corresponding to around 3% of *S. cerevisiae* DNA. Each sample was immunoprecipitated with anti-H3K4me3 (Abcam ab8580) and anti-H3K4me2 (Abcam ab7766) antibodies. Protein A/G beads (Pierce) were coupled to antibodies and resuspended in diluted chromatin and rotated o/n at 4 °C. The next day, beads were washed with RIPA buffer, RIPA-500 buffer (10 mM Tris-HCl pH 8.0, 1 mM EDTA pH 8.0, 0.1% SDS, 500 mM NaCl, 1% Triton X-100, 0.1% sodium deoxycholate), LiCl wash buffer (10 mM Tris-HCl pH 8, 1 mM EDTA, 250 mM LiCl, 0.5% v/v NP-40, 0.5% w/v sodium deoxycholate), and TE buffer (10 mM Tris-HCl pH 8, 1 mM EDTA pH 8). Volume of each wash was 150 µl. Chromatin was eluted from beads in 2×10 µl ChIP elution buffer (Tris-HCl pH 8 50 mM, 1% SDS, 10 mM EDTA pH 8). The immunoprecipitated chromatin as well as a 20 µl sample of input chromatin were decrosslinked by TE, RNase cocktail, Proteinase K and 6 µl SDS (10% w/v), incubating overnight at 65 °C. DNA was purified by ethanol precipitation. Illumina sequencing libraries were prepared from the DNA using the NEBNext Ultra II kit without dual size selection and using 1.4X volume of AMPure XP beads for the purification steps. The libraries were sequenced on Illumina’s NextSeq 500, paired-end, 39 bases from each end.

### Mnase Seq and ChIP Analysis

Illumina adaptor sequences were detected and trimmed using TrimGalore. Trimmed reads were then aligned to Saccharomyces_cerevisiae genome assembly R64-1-1 using BWA. Highly repetitive Ribosome DNA regions were removed from alignment. PCR duplicates were marked and removed by Picard MarkDuplicates. Deduplicted reads from biological replicates were then merged together. Bigwig files were generated from merged bam files by deeptools bamcoverage command with parameters “--binSize 1 -- MNase --minFragmentLength 100 --maxFragmentLength 200 --normalizeUsing CPM”. These parameters aimed to take only mononucleosome, deconvolute and take only the center dyad genomic coordinate of each nucleosome. Bed files containing gene groups generated from RNAseq data were provided to deeptools computematrix command to extract the nucleosome occupancy and average histone modification level of each gene within particular group. Deeptools plotProfile tool was then used to summarise above matrix into metagene plot with parameter “– perGroup”.

### Metabolic labeling and SLAM-Seq

Metabolic labelling of newly synthesized RNA molecules was performed as previously described ^38^. Briefly, 4-thiouracil (4tU) (Sigma) was dissolved in NaOH(83mM), Newly synthesized RNA was labelled for 10 minutes at a final concentration of 5mM 4-tU. MES buffer (pH 5.9) with a final concentration 10mM was added to media to avoid pH change as a result of NaOH addition. At each time points before harvesting (t_0_, t_30_, t_0_’, t_30_’ for both wild type and *rrp6*Δ), prewarmed 4tU was added to culture media (YPD with MES buffer or YPGal with MES buffer) at ten minutes ahead of desired time point. After 10 min labelling with 4tU and reached the desired time point, cells were collected by centrifugation and snap frozen in liquid nitrogen immediately. RNA was purified with MasterPure Yeast RNA Purification Kit. Total RNA was then subjected to thiol(SH)-linked alkylation by iodoacetamide at 50 °C for 15 minutes Seq) ^37^, the reaction was stopped by 20 mM DTT. RNA was cleaned by ethanol precipitation. rRNA was depleted using the RiboPools Depletion Kit (siTOOLs Biotech), strand specific library was prepared by Ultra^™^ II Directional RNA Library Prep Kit for Illumina® following manufacture instruction. Sequencing was performed with single end setting, read length 150bp on Illumina Nextseq 500 sequencer.

Slamseq data was anaysed with slamdunk provided by nextflow pipeline(v 1.0.0).

https://nf-co.re/slamseq. As stranded library was prepared with dUTP method, fastq files were first converted to reverse complementary reads to feed into slamdunk nf core pipeline. Adapter contamination and low-quality region was trimmed using TrimGalore (trim length 30bp). Lifted-over transcript annotation for (Xu et al., 2009) to genome version 64-1-1 was downloaded from SGD database. Annotation was first converted into a bed file and used as input for parameter -utrbed. At least 2 T>C conversions cooccurring in one reads was regarded as a confident call for nascent RNA reads. SNP masking was employed to distinguish Single Nucleotide Polymorphism from converted nucleotides.

