## Supplementary material for "Differential regulation of mRNA stability modulates transcriptional memory and facilitates environmental adaptation"

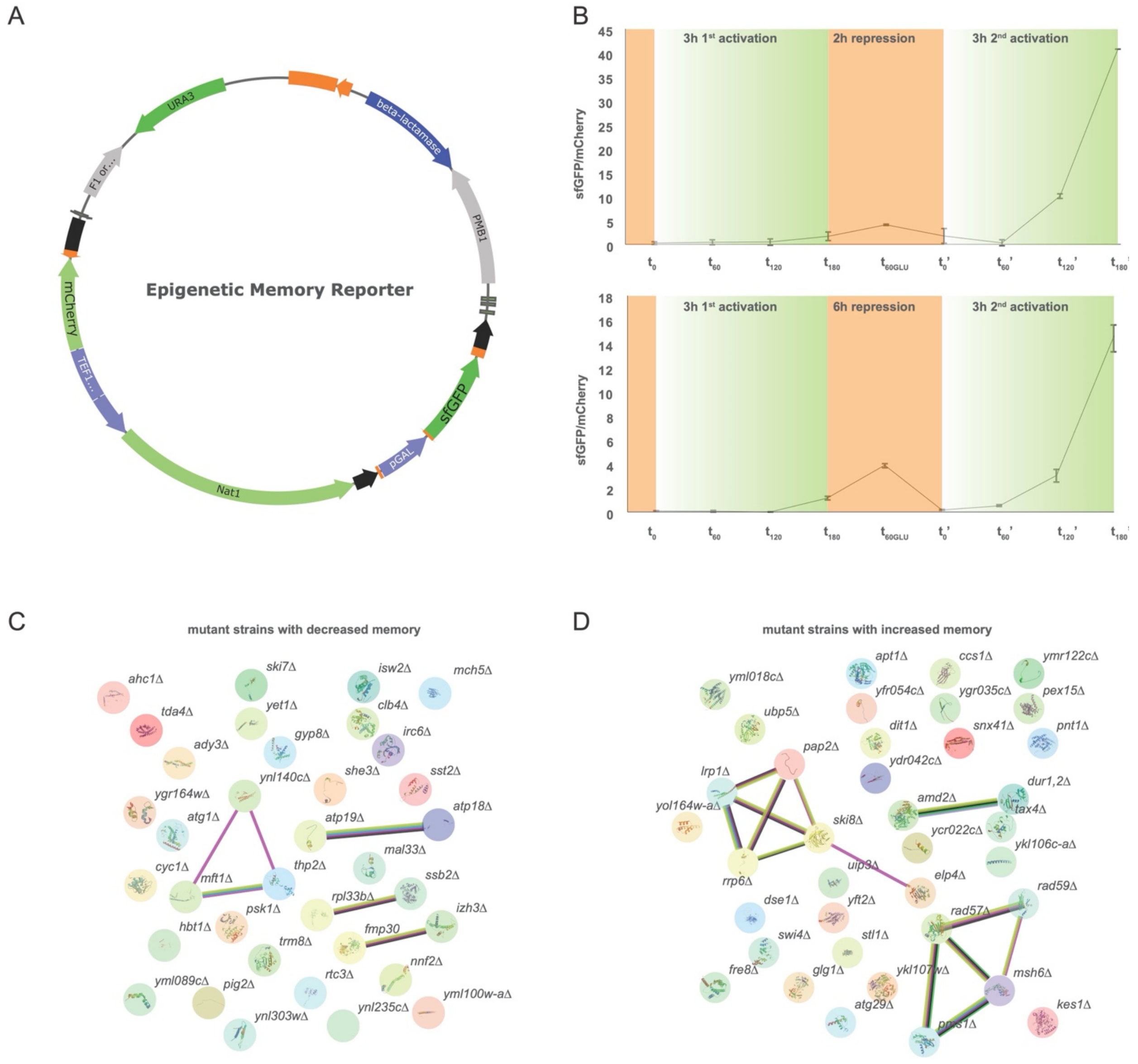


**Fig. S1: Extended information for transcriptional memory screen. A,** Details of the plasmid used as reporter for transcriptional memory using p416 TEF as backbone[41] . **B**, Plate reader measurements of the relative accumulation of *pGAL1-sfGFP* in relation to the constitutively expressed pTEF1-MCherry in a wild-type strain, confirming the ability of the used reporter to display transcriptional memory. Tests using both 2 hours (up) and 6 hours of repression in YPD (down) and 2 transformed clones are shown. As expected, protein accumulation is delayed in respect to transcriptional response. **C**, Network analysis for the 35 candidate genes putatively decreased transcriptional memory (Supplementary Data 1) using STRING v11.5[40]. blue indicate from curated databases, pink indicate experimentally determined, green indicate predicted interaction as gene neighborhood, light green indicate source as text mining, black indicate co-expression. **D**, Network analysis candidate genes putatively enhanced transcriptional memory (Supplementary Data 1) using STRING v11.5[40]. Colour code as in Fig. S1C.

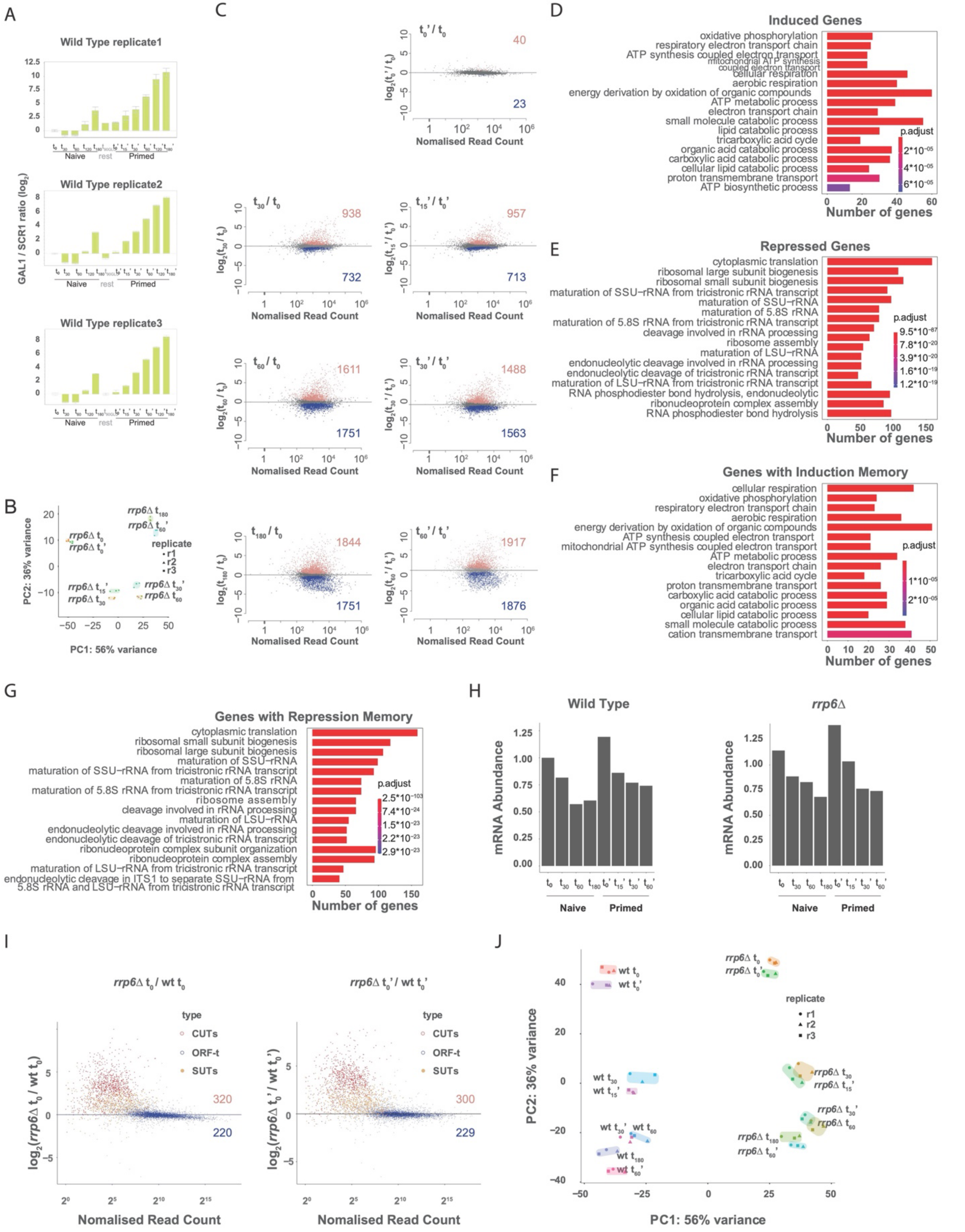
A
Wild Type replicate1
Wild Type replicate2
Wild Type replicate3
GAL1 / SCR1 ratio (log2)
Naive
rest
Primed
B
PC1: 56% variance
PC2: 36% variance
replicate
r1
r2
r3
C
Nomalised Read Count
t0' / t0
40
23
t30 / t0
938
732
t15' / t0'
957
713
t60 / t0
1611
1751
t30' / t0'
1488
1563
t180 / t0
1844
1751
t60' / t0'
1917
1876
D
Induced Genes
oxidative phosphorylation
respiratory electron transport chain
ATP synthesis coupled electron transport
mitochondrial ATP synthesis coupled electron transport
cellular respiration
aerobic respiration
energy derivation by oxidation of organic compounds
ATP metabolic process
electron transport chain
small molecule catabolic process
lipid catabolic process
tricarboxylic acid cycle
organic acid catabolic process
carboxylic acid catabolic process
cellular lipid catabolic process
proton transmembrane transport
ATP biosynthetic process
p.adjust
Number of genes
E
Repressed Genes
cytoplasmic translation
ribosomal large subunit biogenesis
ribosomal small subunit biogenesis
maturation of SSU−rRNA from tricistronic rRNA transcript
maturation of SSU−rRNA
maturation of 5.8S rRNA
maturation of 5.8S rRNA from tricistronic rRNA transcript
cleavage involved in rRNA processing
ribosome assembly
maturation of LSU−rRNA
endonucleolytic cleavage involved in rRNA processing
endonucleolytic cleavage of tricistronic rRNA transcript
maturation of LSU−rRNA from tricistronic rRNA transcript
RNA phosphodiester bond hydrolysis, endonucleolytic
ribonucleoprotein complex assembly
RNA phosphodiester bond hydrolysis
p.adjust
Number of genes
F
Genes with Induction Memory
cellular respiration
oxidative phosphorylation
respiratory electron transport chain
aerobic respiration
energy derivation by oxidation of organic compounds
ATP synthesis coupled electron transport
mitochondrial ATP synthesis coupled electron transport
ATP metabolic process
electron transport chain
tricarboxylic acid cycle
proton transmembrane transport
carboxylic acid catabolic process
organic acid catabolic process
cellular lipid catabolic process
small molecule catabolic process
cation transmembrane transport
p.adjust
Number of genes
G
Genes with Repression Memory
cytoplasmic translation
ribosomal small subunit biogenesis
ribosomal large subunit biogenesis
maturation of SSU−rRNA
maturation of SSU−rRNA from tricistronic rRNA transcript
maturation of 5.8S rRNA
maturation of 5.8S rRNA from tricistronic rRNA transcript
ribosome assembly
cleavage involved in rRNA processing
maturation of LSU−rRNA
endonucleolytic cleavage involved in rRNA processing
endonucleolytic cleavage of tricistronic rRNA transcript
ribonucleoprotein complex subunit organization
ribonucleoprotein complex assembly
maturation of LSU−rRNA from tricistronic rRNA transcript
endonucleolytic cleavage in ITS1 to separate SSU−rRNA from 5.8S rRNA and LSU−rRNA from tricistronic rRNA transcript
p.adjust
Number of genes
H
Wild Type
rrp6Δ
mRNA Abundance
Naive
Primed
I
rrp6Δ t0 / wt t0
rrp6Δ t0' / wt t0'
type
CUTs
ORF-t
SUTs
320
220
300
229
Nomalised Read Count
J
PC1: 56% variance
PC2: 36% variance
replicate
r1
r2
r3

**Fig. S2: Differential regulation of mRNA abundance in naïve and prime states. A,** RT-qPCR analysis of the relative abundance of endogenous GAL1 mRNA normalised to SCR1 for both first and second galactose induction. **B**, Principle component analysis (PCA) of mRNA expression (normalised RNA-Seq data) for both first and second induction in *rrp6*Δ. Only coding mRNAs (ORF-Ts) where considered. **C**, Differential coding gene expression across samples (ORF-Ts). MA plots show $\log_2$ fold change differences on the y-axis and average normalised read counts (by global ORF-Ts abundance) on the x-axis. Time points in naive state are compared to time 0 (naive states). Time points in primed state are compared to time 0' (primed states). Significantly upregulated genes (p-adj < 0.001) are shown in red and significantly down-regulated (p-adj < 0.001) in blue. **D**, Gene Ontology enrichment analysis for induced genes. **E**, Gene Ontology enrichment analysis for repressed genes, **F**, Gene Ontology enrichment analysis for genes with induction memory. **G**, Gene Ontology enrichment analysis for genes with repression memory. **H**, Total RNA abundance (normalised by spike in) of both first and second induction in wild-type and *rrp6*Δ strains. **I**, Differential coding gene expression across samples (ORF-Ts). MA plots show $\log_2$ fold change differences on the y-axis and average normalised read counts (by global ORF-Ts abundance) on the x-axis. Time 0 of *rrp6*Δ in naive state are compared to time 0 of wild type (naive states). Time 0' of *rrp6*Δ in primed state are compared to time 0' of wild type (primed states). **J**, Principle component analysis (PCA) of mRNA expression (normalised RNA-Seq data) for both first and second induction in *rrp6*Δ as in Fig 2B, but including ORF-Ts, CUTs and SUTs. RNA abundance was normalized using coding genes.

A

**Promoter overlap with CUTs**

| | Gene Groups | total | expected | observed | p value | enrich | over |
|---|---|---|---|---|---|---|---|
| Induction Memory | **Attenuated** | 7 | 0.49 | 2 | 0.08 | over | 4.06 |
| | **not changed** | 88 | 6.19 | 10 | 0.09 | over | 1.62 |
| | **enhanced** | 451 | 31.71 | 30 | 0.42 | under | 1.06 |
| | Induced No Memory | 336 | 23.62 | 29 | 0.14 | over | 1.23 |
| | No Change | 3526 | 247.94 | 244 | 0.35 | under | 1.02 |
| | repressed No Memory | 294 | 20.67 | 23 | 0.33 | over | 1.11 |
| Repression Memory | **Attenuated** | 4 | 0.28 | 0 | 0.75 | under | 0 |
| | **not changed** | 611 | 42.96 | 35 | 0.10 | under | 1.23 |
| | **enhanced** | 158 | 11.11 | 12 | 0.43 | over | 1.08 |

C

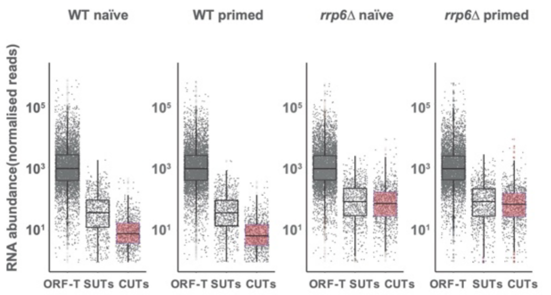


B

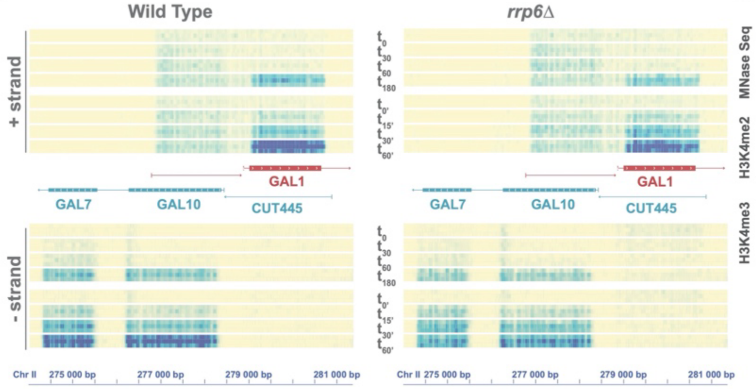


E

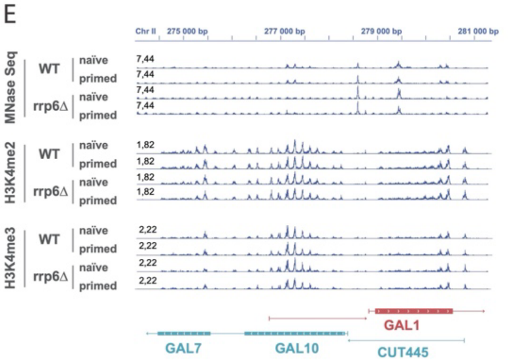


D

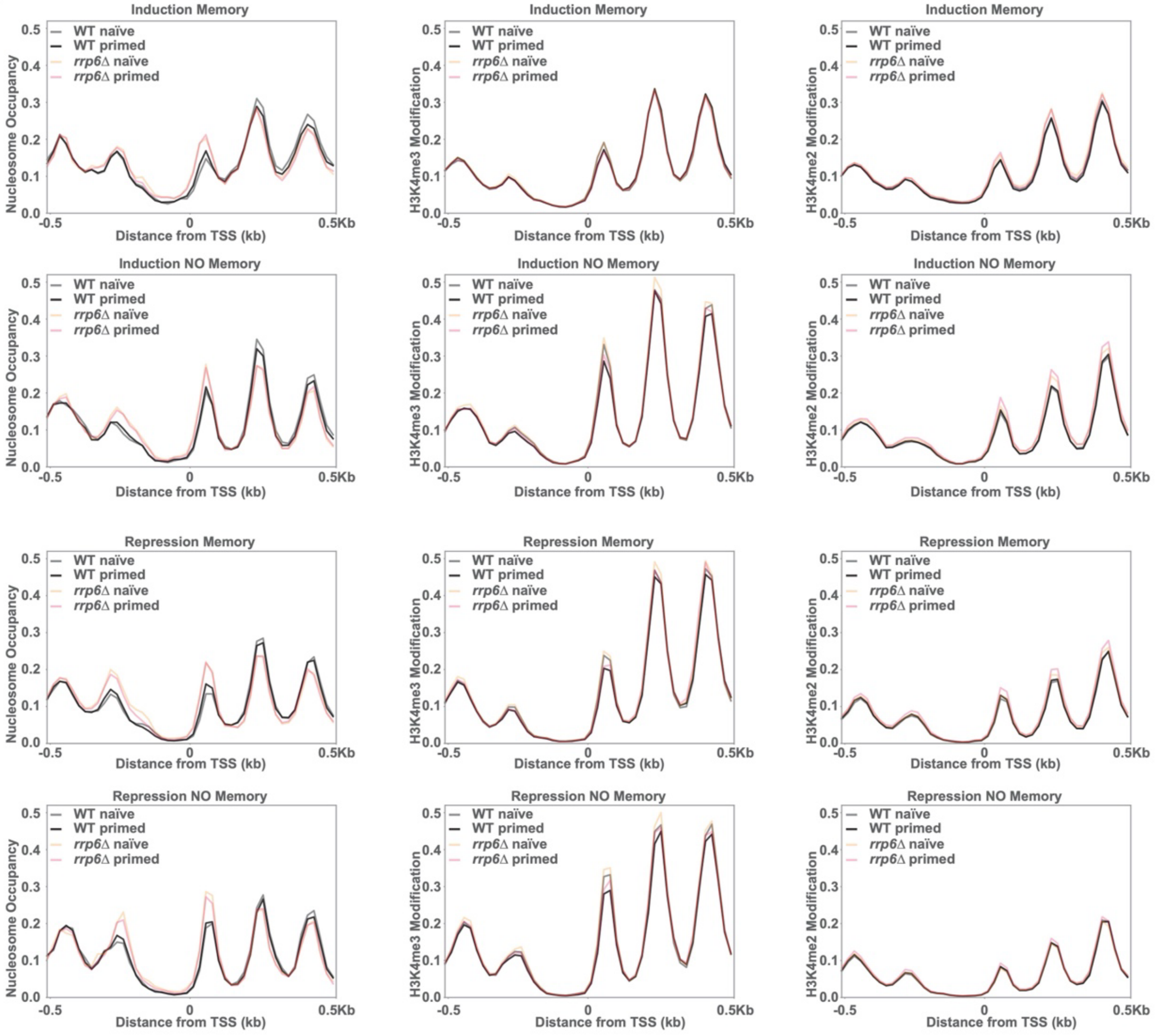

**Fig. S3: Contribution of transcription and chromatin organization to transcriptional memory. A**, Overlap between annotated CUTs and gene groups with different transcriptional memory profiles. Overlap was computed between the transcription start sites and CUTs from[24]. (*i.e.* -150 nt to -50 relative to transcription start site on sense strand and -150 to +50 on antisense strand). Significance for the overlap was tested using a hypergeometric test. **B**, Strand-specific RNA-Seq coverage for the region containing *GAL1*, *GAL10* and *GAL7* in wild type and *rrp6*Δ strains (yellow low expressed to blue, high expressed). **C**, Relative expression of ORF-Ts, SUTS and CUTs in naïve and primed cells for the wild type and the *rrp6*Δ strains. mRNA abundance is normalized to global ORF-Ts abundance. **D**, Global MNase and ChIP Seq analysis. Metaplot of the distribution of average nucleosome, H3K4me3, H3K4me2 signal. Average sequencing coverage is shown (cpm, counts per million) for wild-type $t_0$ naïve (grey), wild-type $t_0$' primed (black), *rrp6*Δ $t_0$ naïve (orange) and *rrp6*Δ $t_0$' primed (pink) here. Genome-wide chromatin profiles around transcription start sites (TSS). Average sequencing coverage is shown (cpm, counts per million) for nucleosome mapping (MNase, left column), H3K4me3 (center column) and H3K4me2 (right column).

FigS4

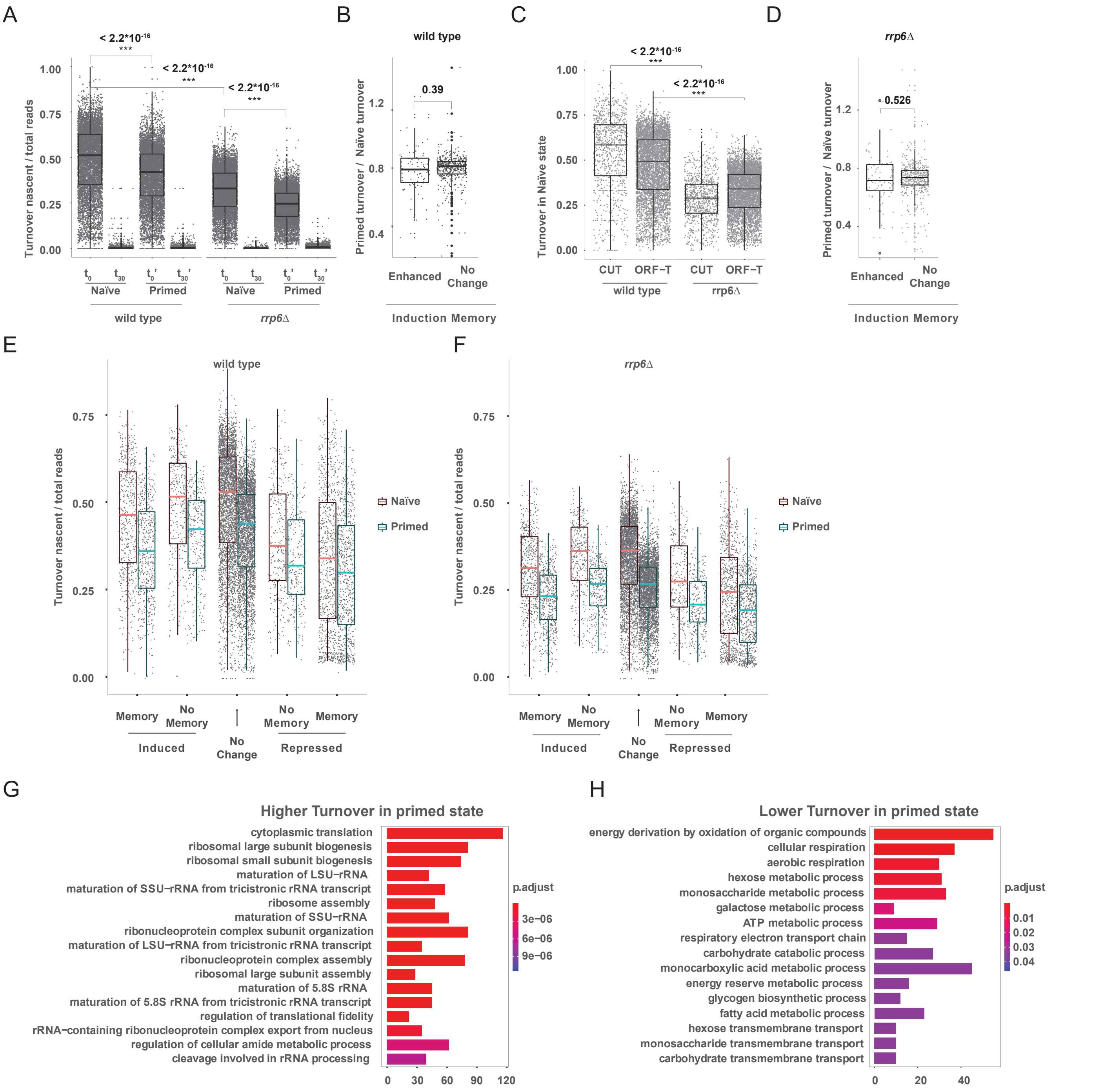


**Fig. S4: Expanded information for SLAM-Seq comparing naïve and primed states.** **A**, Relative mRNA turnover (comparing nascent vs total RNA) using SLAM-seq in naïve and primed conditions at $t_0$ and $t_{30}$ for the wild-type and the *rrp6*Δ strain. **B**-**C,** Gene Ontology enrichment analysis for genes with relatively decreased (B) or increased (C) mRNA turnover in primed cells. **D**, mRNA turnover in naïve ($t_0$) and primed ($t_0$') conditions according to memory gene classification in wild type strain. **E**, in wild type strain, change in mRNA turnover between primed and naïve conditions for genes with induction memory enhanced or not enhanced by *RRP6* depletion. **F**, Relative mRNA turnover (comparing nascent vs total RNA) in wild-type naïve $t_0$ state for CUTs and ORF-Ts. G, As in Fig. S4D but for the *rrp6*Δ strain. **H**, As in Fig. S4E but for the *rrp6*Δ strain.

**Supplementary Data 1: Results of the transcriptional memory screen.** Microsoft Excel workbook containing the summary of cells sorted at each timepoint by FACS, the raw sequencing count for each time window and replicate, and the final list of candidates.

**Supplementary Data 2: Generated RNA-Seq information.** S2A contains raw count, $\log_2$ fold-change, gene type (ORF-T, CUTs or SUTs) and final gene category in column named gene groups. S2A Raw counts are direct output of featurecounts. S2A $\text{Log}_2$ fold change of each sample was compared to time 0 of naïve state in corresponding strain. S2A Gene group column contains nine gene groups: induction memory enhanced in *rrp6*Δ, induction memory that does not change, induction memory attenuated in *rrp6*Δ*,* genes induced without memory, genes with no significant change, genes repressed without memory, repression memory attenuated in *rrp6*Δ, repression memory not changed in *rrp6*Δ and repression memory enhanced in *rrp6*Δ. S2B Comparison of gene expression in naïve state between rrp6Δ and wild type. S2C-S2H Gene Ontology Enrichment for each gene group.

**Supplementary Data 3: Raw data for SLAM-seq analysis. S3A**: Nascent / Total ratio of each gene at time 0 and 30 min in naïve and primed states in both wild type and *rrp6*Δ. GO enrichment of genes which demonstrate relative lower turnover (**S3B**) and higher (**S3C**) turnover in primed state compared to naïve state in wild type.
